# Mechanisms of growth-dependent regulation of the Gin4 kinase

**DOI:** 10.1101/2024.11.20.624605

**Authors:** Francisco Mendez Diaz, David Sanchez Godinez, Francisco Solano, Akshi Jasani, Maria Alcaide, Douglas R. Kellogg

**Affiliations:** Department of Molecular, Cell and Developmental Biology University of California, Santa Cruz

## Abstract

Cell cycle progression is dependent upon cell growth. Cells must therefore translate growth into a proportional signal that can be used to determine when there has been sufficient growth for cell cycle progression. In budding yeast, the protein kinase Gin4 is required for normal control of cell growth and undergoes gradual hyperphosphorylation and activation that are dependent upon growth and proportional to the extent of growth, which suggests that Gin4 could function in mechanisms that measure cell growth. However, the molecular mechanisms that drive hyperphosphorylation of Gin4 are poorly understood. Here, we used biochemical reconstitution and genetic analysis to test hypotheses for the mechanisms that drive phosphorylation of Gin4. We ruled out a previous model in which phosphatidylserine delivered to sites of plasma membrane growth binds Gin4 to initiate autophosphorylation. Instead, we show that Elm1, a homolog of the mammalian Lkb1 tumor suppressor kinase, is sufficient to promote hyperphosphorylation of Gin4 in vitro, likely via initiation of Gin4 autophosphorylation. Furthermore, we show that casein kinase I is required for growth-dependent hyperphosphorylation of Gin4 and also for normal regulation of Elm1. Together, these discoveries lead to new insight into mechanisms that link cell cycle progression to cell growth.

## Introduction

Cell growth is required for cell cycle progression, which indicates that cell growth generates signals that control the cell cycle (Jorgensen and Tyers, 2004; Futcher and Kellogg, 2024) . Maintenance of a specific cell size requires that these growth-dependent signals trigger cell cycle progression only when an appropriate amount of growth has occurred. The mechanisms by which growth-dependent signals are generated, measured, and relayed to the cell cycle machinery are poorly understood.

In both yeast and mammalian cells, growth occurs throughout the cell cycle and is assessed at multiple cell cycle transitions (Liu et al., 2024; Leitao and Kellogg, 2017; Varsano et al., 2017; Soifer et al., 2016; Ferrezuelo et al., 2012). In budding yeast, most cell growth occurs as the daughter bud undergoes rapid expansion during mitosis, which means that maintenance of a specific cell size requires mechanisms that measure and limit bud growth (Leitao and Kellogg, 2017). Bud growth is required for the metaphase to anaphase transition and for mitotic exit, consistent with the idea that bud growth is monitored throughout mitosis (Jasani et al. 2020; Talavera et al. 2023).

How do yeast cells measure and limit growth of the daughter bud to ensure normal control of cell size? There is evidence that the related protein kinases Gin4 and Hsl1 play roles in a mechanism that measures bud growth during early mitosis before metaphase. Loss of Gin4 and Hsl1 leads to a prolonged delay in early mitosis while bud growth continues, leading to formation of aberrantly large buds (Jasani et al., 2020). Thus, loss of Gin4 and Hsl1 causes cells to behave as though they fail to detect that growth has occurred. Furthermore, Gin4 and Hsl1 undergo gradual hyperphosphorylation that is dependent upon bud growth and correlated with the extent of bud growth (Jasani Et al., 2020). Once they have been fully activated, Gin4 and Hsl1 promote progression through mitosis via inactivation of Swe1, the budding yeast homolog of the Wee1 kinase that phosphorylates and inhibits mitotic cyclin/Cdk1 complexes (Longtine et al., 2000; Ma et al., 1996; Jasani et al., 2020). Together, these observations are consistent with the hypothesis that Gin4 and Hsl1 relay growth-dependent signals that control the cell cycle.

Defining the molecular mechanisms that drive hyperphosphorylation and activation of Gin4 and Hsl1 is an essential test of the hypothesis that they relay growth-dependent signals that report on the extent of bud growth. If indeed Gin4 and Hsl1 relay growth-dependent signals, their activation should be mechanistically linked to the events of bud growth. There is already good evidence to support this idea. First, hyperphosphorylation of Gin4 is dependent upon membrane trafficking events that drive plasma membrane growth in the daughter bud (Jasani et al., 2020). Thus, bud growth requires delivery of Golgi-derived vesicles to the plasma membrane, and delivery of these vesicles is also required for hyperphosphorylation of Gin4. The vesicles that drive plasma membrane growth also deliver phosphatidylserine, which becomes highly enriched in the plasma membrane of the growing bud (Moravcevic et al., 2010; Fairn et al., 2011). Gin4 and Hsl1 both have C-terminal KA1 domains that bind phosphatidylserine (Moravcevic et al., 2010; Jasani et al., 2020), and binding of phosphatidylserine to the KA1 domain is necessary for growth-dependent hyperphosphorylation of Gin4 (Jasani et al. 2020). Growth-dependent hyperphosphorylation of Gin4 also requires Gin4 kinase activity, which indicates a role for autophosphorylation (Altman and Kellogg 1997), and mammalian homologs of Gin4 appear to be activated to undergo autophosphorylation by binding to phosphatidylserine (Nesić et al., 2010; Emptage et al., 2017, 2018)). Together, these observations suggested that phosphatidylserine delivered to the growing bud could be both necessary and sufficient to drive growth-dependent phosphorylation of Gin4, which could explain how growth-dependent signals are generated (Jasani et al., 2020). However, this model has never been tested.

Here, we used biochemical reconstitution to test models for the molecular mechanisms that drive growth-dependent phosphorylation of Gin4. The results suggest that binding to phosphatidylserine is not sufficient to drive hyperphosphorylation of Gin4 in vitro, which suggests that phosphatidylserine provides one of several inputs needed for Gin4 hyperphosphorylation. We further discovered that the Elm1 kinase is sufficient to drive Gin4 hyperphosphorylation in vitro. Previous work found that Elm1 is required for Gin4 hyperphosphorylation in vivo. Finally, we found that budding yeast homologs of casein kinase 1, referred to as Yck1 and Yck2, are required for Gin4 hyperphosphorylation in vivo. Since Yck1/2 are membrane-anchored proteins that are delivered to the growing bud via post-Golgi vesicles, they could play an important role in generation of growth-dependent signals.

## Results

### Binding to phosphatidylserine is necessary, but not sufficient, for hyperphosphorylation of Gin4

To carry out in vitro tests of models for the mechanisms that promote Gin4 hyperphosphorylation, we developed a protocol for rapid purification of proteins from yeast that can be carried out at large scale. Briefly, proteins are C-terminally tagged with a tandem affinity tag that includes maltose-binding protein followed by 8XHIS (MBP-8xHIS). A TEV protease cleavage site is included between the protein and the MBP-8XHIS tag to allow removal of the tag. Tagged proteins are bound first to a nickel affinity column followed by an amylose affinity column, which yields highly purified protein. Since both steps are cost effective and easily scale to large volumes of extract, the procedure can be used to purify any protein from yeast. **Figure 1A** shows purification of Gin4-MBP-8XHIS expressed from the endogenous promoter. Previous work found that a protein called Nap1 binds tightly to Gin4 and is required in vivo for normal hyperphosphorylation of Gin4 (Altman and Kellogg 1997). The purified Gin4 is thus in a complex with Nap1.

**Figure 1.**
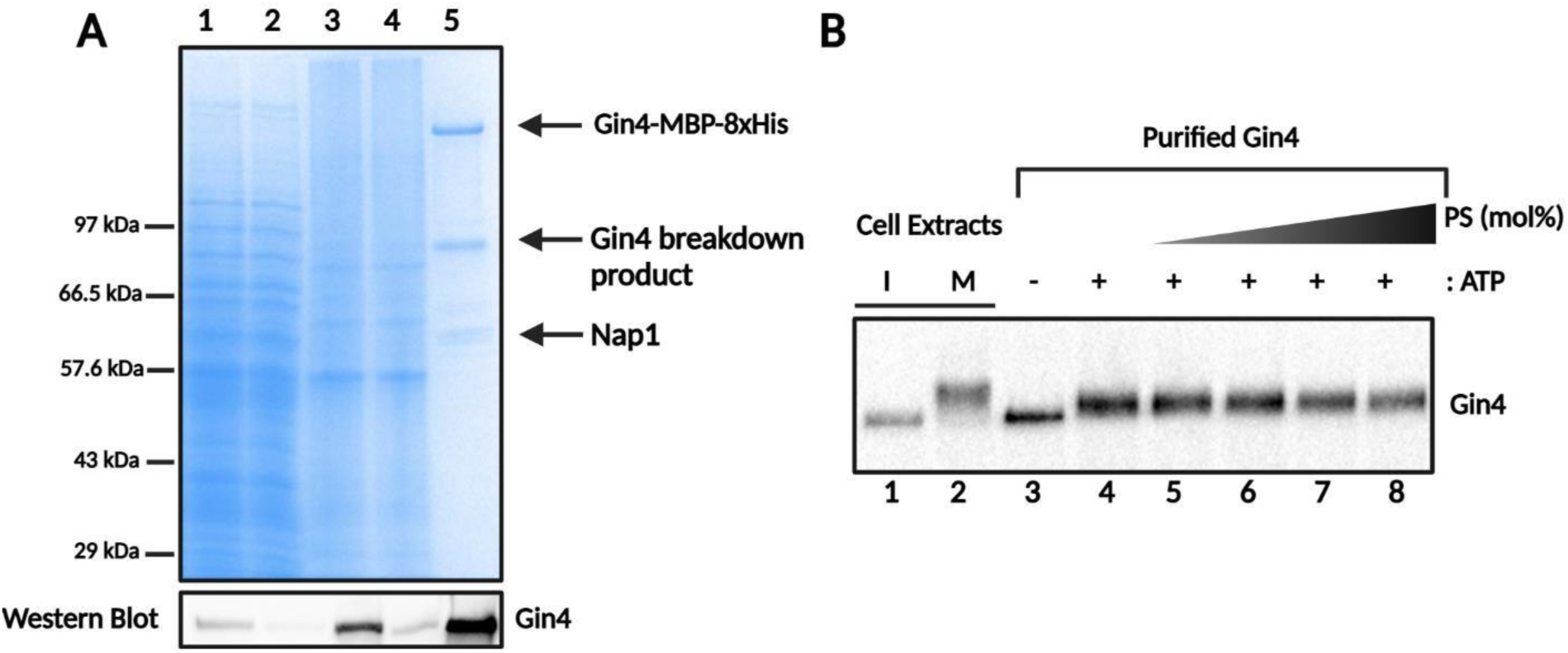
Binding of phosphatidylserine is not sufficient to induce full hyperphosphorylation of Gin4 **A)** Purification of Gin4-MBP-8xHis from log-phase yeast cells. Lane 1: cell extract, lane 2: nickel column flow-through, lane 3: nickel column elution, lane 4: amylose column flow-through, lane 5: amylose column elution. A lower molecular-weight band that is detected with an anti-Gin4 antibody and a likely Gin4 breakdown product is labeled. An anti-Gin4 western blot of the samples shown in the Coomassie blue-stained gel is shown below. **B)** Western blot analysis of Gin4 phosphorylation. Lanes 1 and 2 show the behavior of endogenous Gin4 in extracts from cells arrested in interphase or mitosis. Lanes 3 and 4 show purified Gin4, pre-treated with TEV protease at 30°C for 45 minutes, at 60 nM in the absence and presence of ATP. Lanes 5-9 show the effects of adding phosphatidylserine. Phosphatidylserine was added as lipid vesicles containing phosphatidylcholine and 15%-30% (mol%) phosphatidylserine. The final concentration of phosphatidylserine ranged from 0.075 uM to 0.15 uM. All reactions were carried out at 30°C for 45 minutes.

Since previous studies indicated that autophosphorylation is required for hyperphosphorylation of Gin4, we first tested whether purified Gin4 is capable of undergoing autophosphorylation in the absence of other factors. Hyperphosphorylation of Gin4 was analyzed by western blot to detect electrophoretic mobility shifts caused by phosphorylation. For comparison, we included samples from cells arrested in G1 phase or mitosis. Addition of ATP to Gin4 caused it to undergo autophosphorylation (lanes 3 and 4); however, comparison to Gin4 from mitotic cells indicated that autophosphorylation under these conditions was not sufficient to drive full hyperphosphorylation of Gin4 (**Figure 1B, lanes 1-4)**. These observations indicate that the purified Gin4 has intrinsic kinase activity; however, additional factors are required for full hyperphosphorylation of Gin4.

Previous work with mammalian homologs of Gin4 suggested a model in which binding to phosphatidylserine relieves autoinhibition of Gin4 (Nesić et al., 2010; Emptage et al., 2017, 2018), leading to hyperphosphorylation of Gin4. To test whether binding to phosphatidylserine is sufficient to drive Gin4 hyperphosphorylation, we added lipid vesicles containing phosphatidylcholine and increasing percentages of phosphatidylserine to purified Gin4 in the presence of ATP. Addition of lipid vesicles did not drive further phosphorylation of Gin4 that could be detected via electrophoretic mobility shifts (**Figure 1B, lanes 5-8**). Increasing or decreasing the total amount of added PS-containing vehicles also had no effect on Gin4 phosphorylation. Controls showed that purified Gin4 bound to the phosphatidylserine-containing vesicles in a flotation assay (not shown). The data suggest that binding to phosphatidylserine is not sufficient to induce Gin4 autophosphorylation, but do not rule out the possibility that phosphatidylserine could induce Gin4 hyperphosphorylation under different assay conditions.

### Elm1 can stimulate full hyperphosphorylation of Gin4 in vitro

Previous studies identified candidates for additional factors that directly contribute to hyperphosphorylation of Gin4 (Altman and Kellogg, 1997; Tjandra et al., 1998; Mortensen et al., 2002; Asano et al., 2006; Sreenivasan and Kellogg, 1999). A strong candidate is the protein kinase Elm1. Loss of Elm1 causes a phenotype that is similar to the phenotype caused by loss of Gin4 and Hsl1, and Elm1 is required in vivo for growth-dependent hyperphosphorylation of Gin4 (Sreenivasan and Kellogg, 1999). Elm1 is the budding yeast homolog of the mammalian Lkb1 tumor suppressor kinase and can be functionally replaced by Lkb1, which indicates a high degree of conservation (Sutherland et al., 2003; Hong et al., 2003). Lkb1 kinases phosphorylate a conserved site in the T-loop of AMPK-related kinases, which stimulates their activity (Lizcano et al., 2004; Alessi et al., 2006). Mammalian Lkb1 phosphorylates the MARK kinases, which appear to be homologs of Gin4.

A previous study found that purified Elm1 could promote phosphorylation of Gin4 in vitro (Asano Et al. 2006). However, phosphorylation of Gin4 by Elm1 was only tested with a large molar excess of Elm1 over Gin4, which made it difficult to assess how effectively Elm1 can drive hyperphosphorylation of Gin4. In addition, it was unclear whether Elm1 could drive Gin4 hyperphosphorylation to the same extent that is observed in vivo. We therefore carried out new tests to define how Elm1 contributes to hyperphosphorylation of Gin4.

To obtain purified Elm1, we expressed 6XHIS-GST-Elm1 from the *GAL1* promoter and purified the fusion protein via sequential nickel and glutathione affinity columns, followed by a TEV cleavage step, to obtain highly purified untagged Elm1. The purified Elm1 used for these assays is shown in **Figure 2A**. We carried out a series of reactions with constant amounts of purified Gin4 and increasing amounts of Elm1 and assayed Gin4 hyperphosphorylation by western blot. For reference, we again included samples from cells arrested in interphase and mitosis. This showed that Elm1 is capable of inducing full hyperphosphorylation of Gin4 that matches the extent of hyperphosphorylation observed in vivo for cells arrested in mitosis (**Figure 2B**). Full hyperphosphorylation of 60 nM Gin4 required a minimum of 60 nM Elm1 in a 30-minute reaction at 30°C.

**Figure 2.**
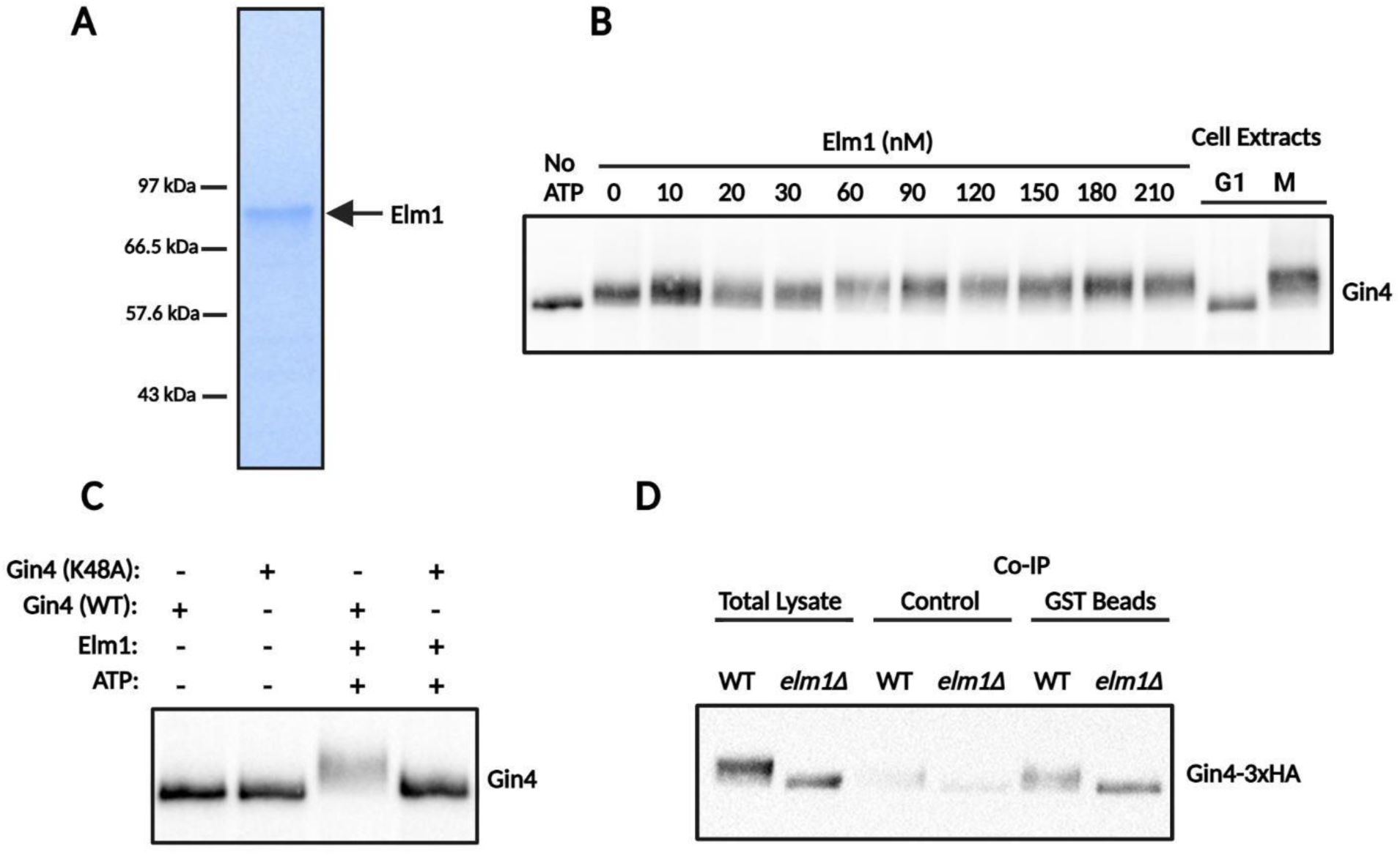
Purified Elm1 can induce full hyperphosphorylation of Gin4 in vitro. **A)** Coomassie blue-stained gel showing the purified Elm1 kinase used for in vitro kinase assays. **B)** Purified Gin4 kinase at 60 nM was pre-cleaved with TEV protease at 30°C to remove the MBP-8XHIS tag and incubated with ATP alone, and with ATP and increasing concentrations of purified Elm1 kinase at 30°C for 45 minutes. Gin4 phosphorylation was assayed by electrophoretic mobility gel shift via western blotting. **C)** Purified wild type and kinase dead Gin4 kinase were incubated with Elm1 kinase in the presence of ATP at 30°C for 45 minutes. Gin4 phosphorylation was assayed by electrophoretic mobility gel shift via western blot. **D)** Cells that contain Gin4-GST and Gin4-3XHA were arrested in mitosis with benomyl and Gin4-GST was immunoprecipitated with anti-GST antibody. As a control, immunoprecipitations were also carried out with a non-specific antibody (anti-MBP). Total lysates and immunoprecipitations were probed with anti- HA to detect Gin4-3XHA.

Elm1 could promote hyperphosphorylation of Gin4 via direct phosphorylation on multiple sites, or it could stimulate Gin4 to undergo autophosphorylation. To distinguish these models, we carried out the same assay with a mutant kinase-dead version of Gin4 purified from yeast. Elm1 was not able to promote hyperphosphorylation of kinase-dead Gin4 (**Figure 2C**). Elm1 and other Lkb1 kinase family members are known to phosphorylate a conserved site within the kinase domain to stimulate kinase activity, whereas data from proteome-wide mass spectrometry analyses have identified over 70 phosphorylation sites that are found outside of the kinase domain and within regions of Gin4 that are predicted to be unstructured. Thus, the data suggest a model in which Elm1 stimulates Gin4 activity to initiate extensive autophosphorylation. However, the data do not rule out an alternative model in which Elm1 induces a limited number of autophosphorylation events that cause Gin4 to undergo conformational changes that promote multi-site phosphorylation by Elm1.

In a previous study, we found that hyperphosphorylation of Gin4 is correlated with initiation of Gin4-Gin4 interactions (Mortensen Et al. 2002). We therefore considered a model in which Elm1 drives Gin4-Gin4 interactions that promote intermolecular autophosphorylation between associated Gin4 molecules. To test this, we used a previously developed assay to determine whether Elm1 is required in vivo for formation of Gin4-Gin4 complexes (Mortensen Et al. 2002). The assay utilizes a strain in which one copy of Gin4 is tagged with 3XHA and another copy is tagged with GST. To test for Gin4-Gin4 interactions, the GST-Gin4 is immunoprecipitated with anti-GST antibodies and western blotting with anti-HA antibody is used to test whether Gin4-3XHA associates with Gin4-GST. We found that Elm1 is not required for Gin4-Gin4 interactions, consistent with a model in which Elm1 stimulates Gin4 kinase activity that is needed for intramolecular Gin4 autophosphorylation (**Figure 2D**).

We also considered a model in which binding of phosphatidylserine promotes phosphorylation of Gin4 by making it a better substrate for Elm1. To test this, we carried out a series of reactions with a low concentration of Elm1 that cannot drive full hyperphosphorylation of Gin4 and then added increasing amounts of lipid vesicles that contain phosphatidylserine. We found no evidence that phosphatidylserine promotes phosphorylation of Gin4 by Elm1 (not shown).

### Yck1/2 kinase activity is required for Gin4 hyperphosphorylation

We next searched for additional factors that could promote hyperphosphorylation of Gin4. Previous studies identified the yeast homologs of casein kinase 1γ as good candidates. Casein kinase 1γ in budding yeast is encoded by a pair of redundant paralogs, referred to as Yck1 and Yck2. Loss of function of Yck1/2 causes defects in control of cell growth that are similar to the phenotype caused by loss of Elm1 or Gin4 and Hsl1 (Lucena et al., 2024; Robinson et al., 1993). Furthermore, inactivation of Yck1/2 with a temperature sensitive allele of YCK2 in a *yck1Δ* background causes a failure to inactivate Swe1 (Pal et al., 2008) . Finally, Yck1/2 are anchored in membranes via a C-terminal palmitoyl group and are delivered to the plasma membrane of the growing bud via membrane trafficking (Babu et al., 2002, 2004). Thus, delivery of Yck1/2 to the growing bud could help generate a growth-dependent signal.

We first tested whether Yck1/2 are required for growth-dependent phosphorylation of Gin4. To do this, we utilized an analog-sensitive allele of *YCK2* in a *yck1Δ* background (*yck2-as yck1Δ*), which allows rapid and specific inhibition of Yck2 activity with the adenine analog 3-MOB-PP1 (Lucena et al., 2024). Wild type control cells and *yck2-as yck1Δ* cells were released from a G1 phase arrest and analog inhibitor was added to both cultures at 15 minutes after release. Hyperphosphorylation of Gin4 was assayed via western blot and levels of the mitotic cyclin Clb2 and Cdk1 inhibitory phosphorylation were assayed as markers of progression through mitosis. Gin4 failed to undergo hyperphosphorylation when Yck1/2 were inhibited, and the cells arrested in mitosis with high levels of Clb2 and sustained Cdk1 inhibitory phosphorylation (**Figures 3)**. Thus, loss of Yck1/2 leads to a failure in growth-dependent phosphorylation of Gin4, as well as a prolonged mitotic delay and a failure to remove Cdk1 inhibitory phosphorylation. Bud growth continues when *yck2-as* is inhibited so the failure in Gin4 phosphorylation is not due to a failure in bud growth.

**Figure 3.**
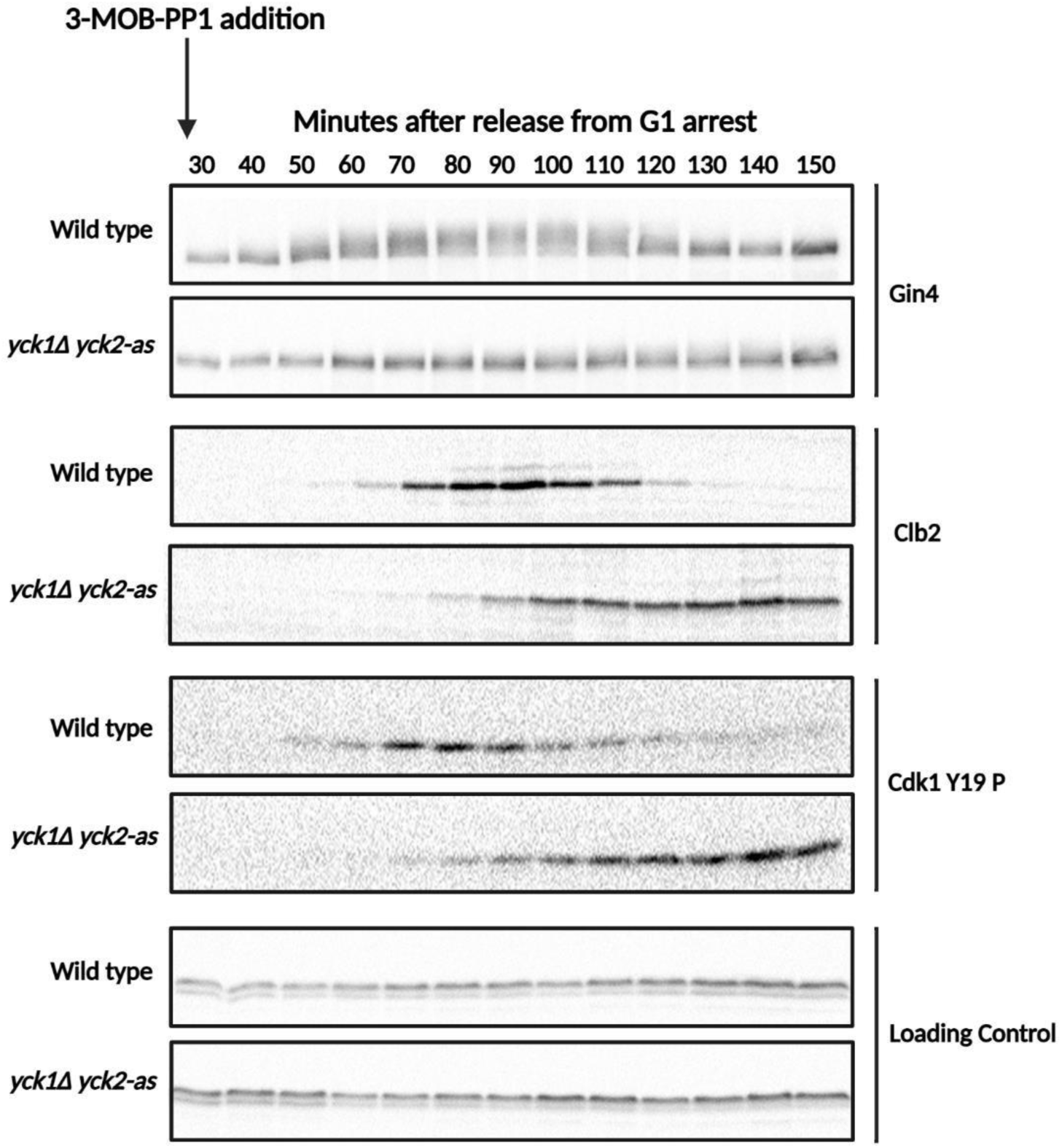
**Yck1/2 kinase activity is required for Gin4 hyperphosphorylation and normal mitotic progression in vivo.** Wild type and *yck1Δ yck2-as* cells were released from a G1 phase arrest in YPD medium without supplemental adenine at room temperature and 3-MOB-PP1 analog was added 15 minutes after release from arrest. Samples were collected at the indicated times and analyzed by western blot. An anti-Nap1 antibody was used for a loading control.

### Localization of Yck1/2 to the plasma membrane is required for Gin4 hyperphosphorylation and normal mitotic progression

Yck1/2 are anchored in the plasma membrane via a C-terminal palmitoyl group that is added by a Golgi-localized palmitoyl transferase referred to as Akr1 (Babu et al., 2002, 2004). Loss of Akr1 causes loss of plasma membrane localization of Yck1/2, as well as defects in control of cell growth. Loss of Yck1/2 is lethal, whereas cells that lack Akr1 are viable, which indicates that Yck1/2 likely have functions that do not require localization to the plasma membrane. To test whether localization of Yck1/2 to the plasma membrane is required for their role in Gin4 hyperphosphorylation, we analyzed Gin4 hyperphosphorylation in *akr1Δ* cells. Wild type and *akr1Δ* cells were released from a G1 phase arrest and Gin4 hyperphosphorylation was assayed by western blot. We also assayed levels of Clb2 and Cdk1 inhibitory phosphorylation as markers of mitotic progression. Hyperphosphorylation of Gin4 failed to occur normally in *akr1Δ* cells compared to wild type cells **(Figure 4)**. Some hyperphosphorylation of Gin4 occurred, likely due to cytoplasmic Yck1/2 that is not associated with the plasma membrane, but a substantial fraction of Gin4 remained in the dephosphorylated form at all time points. Expression of Clb2 and inhibitory phosphorylation of Cdk1 were prolonged **(Figure 4B)**, as expected for a failure to activate Gin4 (**Figure 2C**).

**Figure 4.**
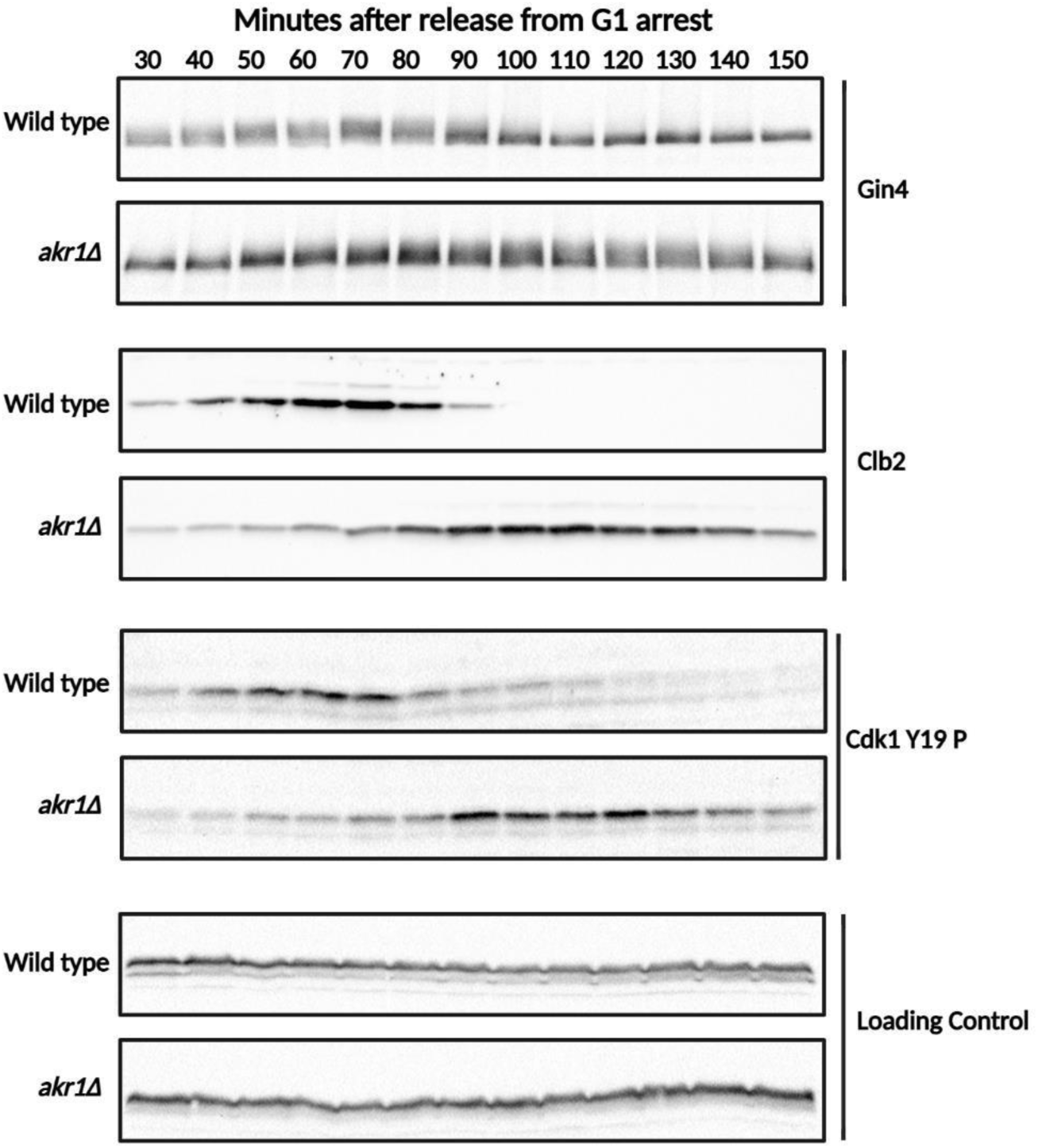
Yck1/2 localization to the plasma membrane is required for Gin4 hyperphosphorylation and normal mitotic progression in vivo. Wild type and *akr1Δ* cells were released from a G1 phase arrest in YPD medium at room temperature. Samples were collected at the indicated times and analyzed by western blot. An anti-Nap1 antibody was used for a loading control.

These observations suggest that Yck1/2 membrane localization is required for normal growth-dependent phosphorylation of Gin4, consistent with a model in which vesicles that drive plasma membrane growth also deliver Yck1/2 to help generate a growth-dependent signal.

### Yck1/2 are required for Elm1 phosphorylation and can directly phosphorylate Elm1 in vitro

We next considered a model in which Yck1/2 directly phosphorylate Gin4. We used purified Yck1 to test the model. The purified Yck1 underwent robust autophosphorylation in vitro, which indicated that it is active. However, we found that Yck1 could not phosphorylate Gin4 in vitro unless it was present at a large molar excess over Gin4, which indicates that it is not capable of efficient phosphorylation of Gin4 (not shown). The data do not rule out models in which additional accessory factors are needed for Yck1/2 to efficiently phosphorylate Gin4. However, in the absence of compelling data that Yck1/2 directly phosphorylate Gin4, we considered a model in which Yck1/2 regulate Elm1. To test this, we determined whether loss of Yck1/2 causes effects on phosphorylation of Elm1. Treatment of Elm1-9xMyc with lambda phosphatase causes it to collapse down to fast migrating forms on a western blot, which indicates that it is a phospho-protein (**Figure 5A**). We next released wild type and *yck2-as yck1Δ* cells from a G1 phase arrest and added analog inhibitor to both cultures at 30 minutes after release from G1 phase arrest. In wild type cells, Elm1 was present in a hyperphosphorylated form and did not undergo large changes in phosphorylation during the cell cycle that could be detected via electrophoretic mobility shifts. In the *yck2-as yck1Δ cells*, Elm1 was not fully in the hyperphosphorylated form before addition of inhibitor and addition of inhibitor caused increased accumulation of dephosphorylated forms of Elm1 (**Figure 5B**). Previous work has shown that the *yck2-as* allele has reduced activity in the absence of inhibitor, which would explain why the *yck2-as* allele causes defects in Elm1 phosphorylation before addition of the inhibitor (Lucena et al., 2024). These results demonstrate that normal hyperphosphorylation of Elm1 requires Yck1/2 activity.

**Figure 5.**
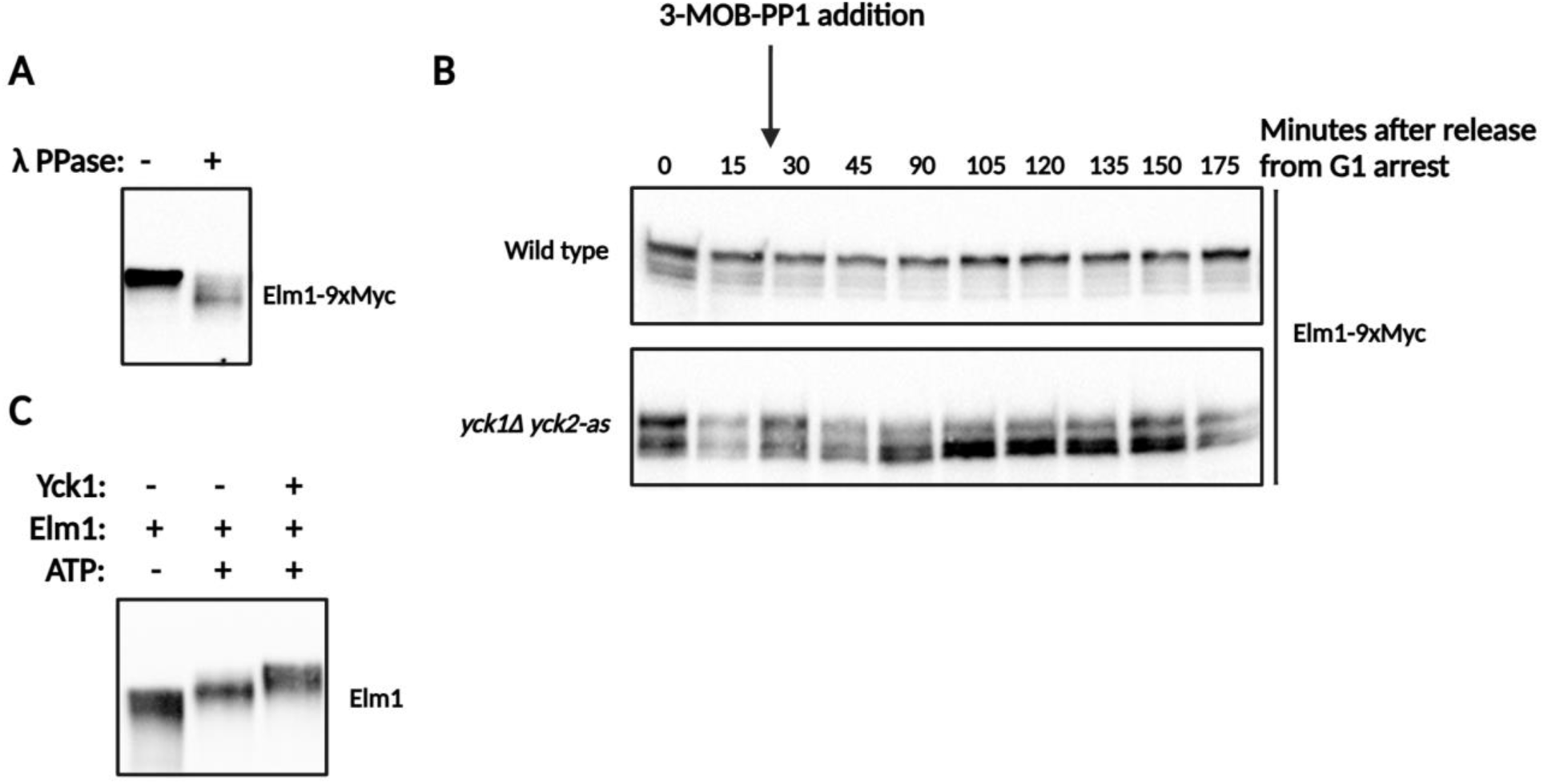
Yck1/2 are required for normal regulation of Elm1. **A)** Elm1-9XMyc cells were lysed in the presence of phosphatase inhibitors or in the presence of lambda phosphatase. Elm1 phosphorylation was assayed by western blot to detect electrophoretic mobility shifts. **B)** Wild type cells and *yck1Δ yck2-as* cells that express Elm1-9XMyc from the endogenous locus were released from a G1 phase arrest into YPD medium without supplemental adenine at room temperature and treated with 3-MOB-PP1 analog 30 minutes after release from the arrest. Samples were collected at the indicated times and analyzed by western blot to detect Elm1-9XMyc. **C)** Purified Elm1 at 200 nM was incubated with ATP alone or with ATP and purified Yck1 at 600 nM at 30°C for 45 minutes. Elm1 phosphorylation was assayed by electrophoretic mobility shift via western blot.

To test whether Yck1/2 can directly phosphorylate Elm1, we determined whether purified Yck1 can phosphorylate Elm1 in vitro. Yck1 was purified from yeast as an MBP-8xHis fusion protein. Elm1 underwent hyperphosphorylation in the presence of ATP (**Figure 5C**). Addition of purified Yck1 caused it to undergo further hyperphosphorylation, which indicates that it is possible that Yck1 directly phosphorylates Elm1 in vivo. A two-fold molar excess of Yck1 was required to achieve full hyperphosphorylation of Elm1 under the conditions used for the reaction. These data show that it is possible that Yck1 phosphorylates Elm1 in vivo, but do not rule out alternative models.

## Discussion

Previous work found that Gin4 is required for normal control of bud growth and undergoes gradual multi-site phosphorylation and activation during bud growth that is dependent upon growth and proportional to the extent of growth, which suggests that Gin4 could play a role in mechanisms that measure growth (Jasani et al. 2020; Altman et al. 1997). Defining the molecular mechanisms that drive hyperphosphorylation of Gin4 is an essential test of the idea that growth generates signals that are relayed via Gin4 to control cell cycle progression. We therefore used biochemical approaches to test models for the mechanisms that drive growth-dependent hyperphosphorylation of Gin4. We found that purified Gin4 undergoes autophosphorylation in the presence of ATP, but the extent of phosphorylation was less than the maximal extent of hyperphosphorylation observed in vivo, which indicates that additional factors are needed. We next tested a model in which hyperphosphorylation of Gin4 is driven by binding to phosphatidylserine that is transported to the plasma membrane during bud growth. This model was suggested by our previous discovery that activation of Gin4 in vivo requires binding of phosphatidylserine to the KA1 domain (Jasani et al. 2020), and that binding of phosphatidylserine to the KA1 domain of related mammalian MARK kinases appears to relieve autoinhibition of kinase activity (Nesic et al. 2010; Emptage et al. 2017). We found no evidence that binding to phosphatidylserine is sufficient for Gin4 hyperphosphorylation in vitro. We therefore favor a model in which binding to phosphatidylserine is required for recruiting Gin4 to the plasma membrane where additional factors drive Gin4 hyperphosphorylation, or a model in which phosphatidylserine binding causes a conformational change in Gin4 that makes it competent to undergo a currently unknown but essential step leading to hyperphosphorylation.

We next tested how additional kinases contribute to hyperphosphorylation of Gin4. We found that the Elm1 kinase is sufficient to drive full hyperphosphorylation of Gin4 in vitro, and that Elm1-dependent hyperphosphorylation of Gin4 requires Gin4 kinase activity, which indicates that autophosphorylation events are involved. Elm1 is a member of a conserved family of Lkb1-related kinases that stimulate the activity of AMPK-related kinases, including Gin4, via phosphorylation of a conserved site in the kinase activation loop (Lizcano et al., 2004; Alessi et al., 2006). Thus, the data suggest that Elm1-dependent phosphorylation of the Gin4 kinase domain likely stimulates Gin4 to undergo extensive autophosphorylation. This raises the question of how Elm1 contributes to gradual growth-dependent hyperphosphorylation of Gin4. We considered a model in which a growth-dependent signal drives a gradual increase in Elm1 kinase activity, which drives gradually increasing activation and autophosphorylation of Gin4. In this model, one might expect to see Elm1 undergo changes in phosphorylation as it becomes activated that are correlated with Gin4 hyperphosphorylation. We did not detect substantial changes in Elm1 electrophoretic mobility in wild type cells that would indicate major changes in phosphorylation; however, not all phosphorylation events cause changes in electrophoretic mobility, so it is possible that Elm1 undergoes phosphorylation changes that are correlated with Gin4 hyperphosphorylation but not detectable via electrophoretic mobility shifts. An alternative model is that Elm1 is necessary to initiate extensive Gin4 autophosphorylation events, but additional factors enforce gradual hyperphosphorylation of Gin4 in vivo. In this model, Elm1 would provide a permissive signal for Gin4 hyperphosphorylation, but other factors would enforce gradual growth-dependent hyperphosphorylation.

We used a candidate approach to search for additional factors that influence growth-dependent hyperphosphorylation of Gin4. The Yck1/2 kinase paralogs were strong candidates because loss of function mutants cause defects in control of cell growth and size that look similar to the defects caused by loss of Gin4 and Hsl1 (Lucena et al., 2024; Robinson et al., 1993). Moreover, Yck1/2 are membrane-anchored proteins that are transported to the plasma membrane of the growing bud via the vesicles that drive bud growth (Babu et al., 2002, 2004). We found that Yck1/2 are required for growth-dependent phosphorylation of Gin4, and that localization of Yck1/2 to the plasma membrane is required for normal growth-dependent phosphorylation of Gin4. These results suggest that the vesicles that drive plasma membrane growth deliver both phosphatidylserine and Yck1/2 to promote growth-dependent phosphorylation of Gin4. We did not find compelling evidence that Yck1/2 are capable of efficiently phosphorylating Gin4 in vitro, which suggests that they promote hyperphosphorylation of Gin4 indirectly. Together, the data provide new insight into the molecular mechanisms responsible for growth-dependent phosphorylation of Gin4 and suggest new models for how growth generates a signal that promotes progression through mitosis. It appears that Elm1-dependent phosphorylation of the Gin4 kinase domain stimulates Gin4 kinase activity to initiate extensive autophosphorylation of Gin4. Autophosphorylation would be expected to be a rapid intramolecular reaction, yet the extent of Gin4 phosphorylation gradually increases during bud growth in vivo, which suggests the existence of mechanisms that restrain autophosphorylation. A model that could explain the data is that autophosphorylation of Gin4 is opposed by a phosphatase that must be inhibited to achieve full hyperphosphorylation of Gin4. In this case, delivery of Yck1/2 to the growing plasma membrane could initiate inhibition of the phosphatase, and the extent of phosphatase inhibition would gradually increase as more and more Yck1/2 are delivered to the growing membrane, leading to gradually increasing Gin4 hyperphosphorylation. Loss of Yck1/2 would lead to hyperactivity of the phosphatase, which would prevent Gin4 hyperphosphorylation. Hyperactivity of a phosphatase could also explain why hyperphosphorylation of Elm1 is reduced in *yck2-as* cells. Identification and characterization of phosphatases that work on Gin4 will be an important next step towards further testing of models for how growth-dependent hyperphosphorylation of Gin4 is achieved.

## Materials and Methods

### Yeast strain construction, plasmid construction, media, and reagents

All yeast strains are in the W303 background (leu2-3 112 ura3-1 can1-100 ade2-1 his3-11, 14 trp1- 1, GAL+, ssd1-d2). Additional genetic features are listed in Table 1. Yeast cells were cultured in YP medium (yeast extract, peptone, 40 mg/L adenine) supplemented with 2% dextrose (YPD), 2% galactose (YPGal), or 2% glycerol/2% ethanol (YPG/E), as indicated in the text. In experiments involving analog-sensitive alleles of kinases, cells were grown in media that was not supplemented with adenine.

**Table 1:** Strains used in this study.

| Strain | Genotype | Citation | Figure |
| --- | --- | --- | --- |
| DK186 | *MATa bar1* |  | Figure 3 and 4 |
| DK4657 | *MATa GIN4-MBP-8xHIS::HIS3 bar1* | This study | Figure 1 |
| DK4705 | *MATa kanMX6::GAL1pr-GIN4-MBP-8xHIS::HIS3,GAL1::kanMX6 bar1* | This study | Figure 1 and 2 |
| DK4712 | *MATa gin4<sup>K48A</sup>-MBP-8xHIS::HIS3 bar1* | This study | Figure 2 |
| EM9 | *MATa URA3::GIN4-GST, TRP1::GIN4-3XHATRP1 (pEM103) bar1* | Mortensen et al. | Figure 2 |
| DK4329 | *MATa URA3::GIN4-GST, TRP1::GIN4-3XHATRP1 (pEM103) elm1Δ bar1* | This study | Figure 2 |
| DK1391 | *MATa yck1Δ::kanMX6 yck2-as bar1* | This study | Figure 3 |
| DK1395 | *MATa akr1Δ::TRP1* | This study | Figure 4 |
| DK177 | *MATa* |  | Figure 4 |
| DK4818 | *MATa ELM1-9xMYC::hphNT1 bar1* | This study | Figure 5 |
| DK4807 | *MATa yck1Δ::kanMX6, yck2-as ELM1-9xMYC::hphNT1 bar1* | This study | Figure 5 |
| DK4434 | *MATa/α URA3::GAL1pr-6xHIS-GST-ELM1, GAL10pr-GAL4 (pBP1) bar1* | This study | Figure 2 and 5 |
| DK4751 | *MATa kanMX6::GAL1pr-YCK1-MBP-8xHIS::HIS3 bar1* | This study | Figure 5 |

Gene deletion, C-terminal tagging, and integration of the *GAL1* promoter in front of genes was carried out by standard PCR-based homologous recombination (Longtine et al., 1998; Janke et al., 2004). To add MBP-8XHIS tags to proteins, a plasmid was created that allows rapid PCR-based tagging. Briefly, yeast codon-optimized MBP-8XHIS DNA sequence with flanking PacI and AscI restriction sites was synthesized (TWIST Biosciences), amplified by PCR, and used to replace the GST tag in pFA6a-GST- HIS3MX6 (Longtine et al., 1998), and verified by DNA sequencing *(pFM26*). To create a plasmid for high level expression of 6xHis-GST-TEV-ELM1, the Elm1 open reading frame, codon-optimized for high level expression, was synthesized and cloned into the HindIII and BamH1 sites of pDK125 to create pBP1. pDK125 is an integrating vector that expresses proteins that are N-terminally tagged with codon-optimized 6XHIS-GST-TEV from the *GAL1* promoter, and also expresses *GAL4* from the *GAL10* promoter, which drives higher level expression from the *GAL1* promoter.

Yeast extract and peptone powder were purchased from BP Diagnostics. Amylose resin for purification of MBP-tagged proteins was purchased from New England BioLabs. Nickel resin for purification of His-tagged proteins was purchased from Sigma-Aldrich (cOmplete^TM^ His-Tag resin). All lipids, including brain PS, egg PC, and DGS (Ni) were purchased from Avanti Polar Lipids, Inc.

### Cell cycle time course experiments and western blotting

Cell cycle time course experiments were conducted as previously described (Harvey et al., 2011). For the experiment in Figure 3, wild type and *yck1Δ yck2-as* yeast cells were grown overnight in YPD medium without supplemental adenine at room temperature to an optical density (OD_600_) between 0.4-0.7. Wild type cells were adjusted to an OD_600_ of 0.5 and *yck1Δ yck2-as* cells were adjusted to 0.65. Cells were arrested in G1 phase by addition of alpha factor to a final concentration of 0.5 µg/mL for three hours at room temperature. Cells were released from the arrest by three washes with medium without alpha factor and incubated at 25°C with gentle agitation. Alpha factor was added back to cultures 60 minutes after release to prevent initiation of a second cell cycle. 3-MOB-PP1 inhibitor was added to 5 uM to both cultures at time point 15 minutes. At each time point, 1.6 mL of sample were collected in screw cap tubes and centrifuged at 15,000 rpm for 15 seconds. The supernatant was then removed and 200 uL of acid- washed glass beads were added before freezing samples in liquid nitrogen. Samples were processed for SDS-PAGE gel electrophoresis on a 10% polyacrylamide gel and western blot as previously described (Harvey et al., 2011).

For the time course data in Figure 4, wild type and *akr1Δ* yeast cells were grown in YPD medium supplemented with adenine and synchronized with 15 ug/mL alpha factor. A higher concentration of alpha factor was used because the strains are *BAR1+*. Cell samples were collected and processed for western blot analysis as described above.

Blots were probed with primary antibody overnight at 4°C in western wash buffer (1x phosphate-buffered saline, 250 mM NaCl, and 0.1% Tween-20) containing 5% w/v nonfat dry milk. Primary antibodies to detect Gin4, Nap1, and Clb2 were rabbit polyclonal antibodies generated as described previously and used at 1-2 ug/ml (Kellogg and Murray, 1995; Altman and Kellogg, 1997; Mortensen et al., 2002) Cdk1 phosphorylated at tyrosine 19 was detected using a Cdc2 Y15 monoclonal primary antibody (Cell Signaling Inc. #9111). Primary antibody to detect Myc-tagged Elm1 was a monoclonal antibody (Cell signaling Inc. #9402). Primary antibodies were detected with HRP-conjugated donkey anti-rabbit secondary antibody (for Gin4, Nap1, and Clb2) or HRP-conjugated donkey anti-mouse secondary antibody (Cdk1-Y19P, Myc) incubated in 5% milk western wash buffer for 1 hour at room temperature. Blots were rinsed in western wash buffer, followed by a final wash in 1x PBS before detection via chemiluminescence and a Bio-rad ChemiDoc imaging system.

### Purification of MBP-8xHis tagged yeast proteins

Cells that express full length MBP-8xHis-tagged Gin4 and Yck1 from the *GAL1* promoter were grown in 6 L of YP medium supplemented with 2% glycerol/2% ethanol for 12-16 hours at 22°C until log phase (OD_600_ 0.4-0.6) was reached. Galactose powder was added to each culture to a 2% final (w/v) concentration and cultures were transferred to 30°C for an additional 4 hours. Cells were harvested by centrifugation at 5,000 rpm at 4°C using a JLA 9.1000 Beckman rotor. Cell pellets were scooped out using a spatula and immediately flash-frozen in liquid nitrogen. Lysis was performed by first grinding frozen chunks of cells in dry ice using a coffee grinder for 30 seconds, followed by further grinding for 15-20 minutes in a mortar and pestle pre-chilled with liquid nitrogen. The resulting powder was stored for 12- 16 hours at -80°C to allow for sublimation of carbon dioxide from the dry ice used in the first grinding step.

To create a cell lysate, 10-15 grams of cell powder was resuspended in 3 volumes of ice-cold lysis buffer (50 mM HEPES-KOH, pH 7.6, 175 mM KCl, 0.5% Tween-20, 1 mM MgCl2, 1 mM EGTA, 10 mM Beta-glycerol phosphate, and 2 mM PMSF). In Yck1 protein purifications, no beta-glycerol phosphate was used in the lysis buffer. The crude lysate was centrifuged at 20,000 RPM for 20 minutes at 4°C and the supernatant was applied to a pre-equilibrated 5 mL nickel column (cOmplete nickel beads, Sigma-Aldrich) at 20 mL/hour using a peristaltic pump at 4°C. Use of the cOmplete nickel beads, rather than Ni-NTA beads, allowed addition of EDTA to the purification buffers, which may help limit proteolysis. The column was then washed with 10 column volumes of nickel column washing buffer (50 mM HEPES-KOH, pH 7.6, 175 mM KCl, 5 mM Imidazole, 1 mM EGTA, 1 mM DTT) at 50 mL/hour. MBP-8xHis tagged proteins were eluted from the column by applying 1.5 mL fractions of nickel elution buffer (50 mM HEPES-KOH pH 7.6, 175 mM KCl, 100 mM Imidazole, 1 mM EGTA, 1 mM DTT, 10% glycerol). Eluted protein was detected via Bradford assay and immediately applied to a 1 mL amylose column at 4°C (New England Biolabs) at 10-15 ml/hr that was pre-equilibrated with nickel elution buffer. For the Yck1 purification, an additional washing step with 5 column volumes of buffer containing 175 mM KCl was performed to reduce salt concentration. Proteins were eluted with 1 mL of MBP elution buffer (50 mM HEPES-KOH pH 7.6, 175 mM KCl, 1 mM DTT, 10 mM Maltose, and 10% glycerol). Eluted proteins were supplemented with MgCl2 to a final concentration of 1 mM and Tween-20 to a final concentration of 0.01%. The concentration of proteins was estimated by running a sample on SDS-PAGE, staining with Coomassie blue, and comparing the band intensity to a standard BSA curve ranging from 0.1-3 ug. The estimated yield of MBP-8xHis-Gin4 was 250 ug and MBP-8xHis-Yck1 was 3000 ug from 10-15 g of cells. Kinase-dead Gin4 expressed from the endogenous promoter was purified as an MBP-8xHis fusion protein. Purified proteins were aliquoted, flash-frozen in liquid nitrogen, and stored at -80°C for in vitro kinase assays.

### Purification of Elm1

Cells that express 6xHis-GST-ELM1 from the *GAL1* promoter were grown in YP galactose medium and harvested and lysed as described for purification of 8XHIS-MBP fusions. 12 grams of cell powder were resuspended in 3 volumes of ice-cold lysis buffer (50 mM HEPES-KOH pH 7.6, 1 M KCl, 0.5% Tween-20, 20 mM imidazole, 2 mM EDTA, 75 mM Beta-glycerol phosphate, 10% glycerol and 2 mM PMSF) and the crude extract was centrifuged at 20,000 RPM at 4°C. The supernatant was applied to a 5 mL nickel column (cOmplete nickel beads, Sigma-Aldrich) at 20 mL/hour at 4°C. After sample application, the column was washed with 40 column volumes of wash buffer (50 mM HEPES-KOH pH 7.6, 1 M KCl, 20 mM imidazole, 2 mM EDTA, and 10% glycerol) overnight at 4°C to remove any endogenous glutathione from the crude extract that could remain bound to GST. The 6xHis-tagged Elm1 protein was eluted from the nickel column by applying 1.5 mL fractions of nickel elution buffer (50 mM HEPES-KOH pH 7.6, 1 M KCl, 300 mM imidazole, 2 mM EDTA, and 10% glycerol). Peak fractions were detected via Bradford assay and then immediately applied to a 1 mL glutathione-agarose column at 10 mL/hour at 4°C that was pre-equilibrated with elution buffer. The glutathione column was then washed with 15 column volumes of glutathione wash buffer (50 mM HEPES-KOH pH 7.6, 300 mM KCl, 1 mM DTT, 1 mM EDTA, and 10% glycerol) and eluted in 150 uL fractions with glutathione elution buffer (50 mM HEPES-KOH pH 7.6, 150 mM KCl, 1 mM DTT, 10 mM reduced glutathione, 10% glycerol). Peak fractions were detected via Bradford assay and pooled. The 6xHis-GST tag was removed by TEV cleavage overnight at 4°C, followed by running the cleavage reaction over two sequential 100 uL nickel columns. The concentration was estimated as described for 8XHIS-MBP fusions. The estimated yield of purified Elm1 was 1000 ug from 12 g of cells. Purified Elm1 was aliquoted, flash-frozen in liquid nitrogen, and stored at -80°C for *in vitro* kinase assays.

### Preparation of liposomes for *in vitro* kinase assays

Lipid suspensions of 15% PS/ 85% PC, 20% PS/80% PC, 25% PS/75% PC, and 30% PS/70% PC (mol%) were made in low-retention microcentrifuge tubes and dried in a speed vac at high temperature for 20 minutes. The lipid film was then washed with 50 µL of sterile water and dried again. Lipids were then resuspended in liposome storage buffer (20 mM Tris-HCl, pH 7.8, 50 mM potassium acetate, 1 mM magnesium acetate) and incubated at 60°C in a water bath for 30 minutes. Six freeze/thaw cycles were then performed: Samples were frozen in liquid nitrogen and thawed at 60°C in a water bath for 5 minutes. Lipid suspension was then passed through an Avanti extruder pre-heated to 60°C for a total of 8-10 passes. Liposomes were stored at 4°C for a maximum of 5 days. The final concentration of lipids used for liposome preparation was 1 mM and they were used at 0.5 µM final concentration in in vitro kinase reactions. In assays where we decreased or increased the total amount of added PS-containing vesicles, the total lipid concentration ranged from 0.1 uM to 1.5 uM.

### Kinase assays

Before addition to assays, purified MBP-8xHis-tagged proteins were pre-treated with His-tagged TEV protease for 2.5 hours at 30°C to remove the tag. In all experiments, Gin4 kinase was used at 60 nM final concentration. Purified Elm1 was used at a range of 10-210 nM in the experiment described in Figure 2B. Elm1 was used at 60 nM in the experiment described in figure 2C. For all other in vitro kinase assays, Elm1 was used at 20 nM final concentration and Yck1 was used at 400 nM final concentration (Figure 5). Purified Gin4 kinase was reconstituted in varying conditions with liposomes, Elm1, and Yck1 in kinase assay buffer (20 mM Tris-acetate pH 7.8, 50 mM Potassium acetate, 1 mM Magnesium acetate, 0.5 ug BSA, 1 mM MnCl2, and 1 mM ATP) in a final volume of 25 µL. In vitro reactions were incubated at 30°C for 30 minutes and quenched by adding 2x sample buffer and incubating at 95°C for 5 minutes. Gin4 kinase was in a dephosphorylated state as shown in Figure 1, therefore, it was not necessary to treat Gin4 with lambda phosphatase prior to the in vitro kinase assay.

Purified Elm1 kinase was reconstituted in varying conditions with Yck1 in kinase assay buffer (20 mM Tris-acetate pH 7.8, 50 mM potassium acetate, 1 mM magnesium acetate, 0.5 ug BSA, 1 mM MnCl2, and 1 mM ATP) for a final volume of 25 µL. In vitro reactions were incubated at 30 ◦C for 45 minutes, then quenched by adding 2x sample buffer and boiling of the samples at 95 for 5 minutes. Elm1 phosphorylation was assayed by electrophoretic mobility gel shift via western blotting, as described above. Elm1 was not treated with lambda phosphatase prior to the in vitro kinase assay.

### Gin4 dimerization assay and treatment of Elm1 with phosphatase

Tests for Gin4-Gin4 interactions and treatment of Elm1 with phosphatase were carried out as previously described (Mortensen et al., 2002; Lucena et al., 2017).

## Acknowledgements

This work was supported by NIH Grant R35 GM131826.

